# SLIDR and SLOPPR: Flexible identification of spliced leader *trans*-splicing and prediction of eukaryotic operons from RNA-Seq data

**DOI:** 10.1101/2020.12.23.423594

**Authors:** Marius A. Wenzel, Berndt Müller, Jonathan Pettitt

## Abstract

**Background:** Spliced leader (SL) *trans*-splicing replaces the 5’ end of pre-mRNAs with the spliced leader, an exon derived from a specialised non-coding RNA originating from elsewhere in the genome. This process is essential for resolving polycistronic pre-mRNAs produced by eukaryotic operons into monocistronic transcripts. SL *trans*-splicing and operons may have independently evolved multiple times throughout Eukarya, yet our understanding of these phenomena is limited to only a few well-characterised organisms, most notably *C. elegans* and trypanosomes. The primary barrier to systematic discovery and characterisation of SL *trans*-splicing and operons is the lack of computational tools for exploiting the surge of transcriptomic and genomic resources for a wide range of eukaryotes.

**Results:** Here we present two novel pipelines that automate the discovery of SLs and the prediction of operons in eukaryotic genomes from RNA-Seq data. SLIDR assembles putative SLs from 5’ read tails present after read alignment to a reference genome or transcriptome, which are then verified by interrogating corresponding SL RNA genes for sequence motifs expected in *bona fide* SL RNA molecules. SLOPPR identifies RNA-Seq reads that contain a given 5’ SL sequence, quantifies genomewide SL *trans*-splicing events and predicts operons via distinct patterns of SL *trans*-splicing events across adjacent genes. We tested both pipelines with organisms known to carry out SL *trans*-splicing and organise their genes into operons, and demonstrate that 1) SLIDR correctly detects expected SLs and often discovers novel SL variants; 2) SLOPPR correctly identifies functionally specialised SLs, correctly predicts known operons and detects plausible novel operons.

**Conclusions:** SLIDR and SLOPPR are flexible tools that will accelerate research into the evolutionary dynamics of SL *trans*-splicing and operons throughout Eukarya and improve gene discovery and annotation for a wide-range of eukaryotic genomes. Both pipelines are implemented in Bash and R and are built upon readily available software commonly installed on most bioinformatics servers. Biological insight can be gleaned even from sparse, low-coverage datasets, implying that an untapped wealth of information can be derived from existing RNA-Seq datasets as well as from novel full-isoform sequencing protocols as they become more widely available.

## Background

Spliced leader (SL) *trans*-splicing is a eukaryotic post-transcriptional RNA modification whereby the 5’ end of a pre-mRNA receives a short “leader” exon from a non-coding RNA molecule that originates from elsewhere in the genome [1, 2]. This mechanism was first discovered in trypanosomes [3] and has received much attention as a potential target for diagnosis and control of a range of medically and agriculturally important pathogens [1, 4, 5]. SL *trans*-splicing is broadly distributed among many eukaryotic groups, for example euglenozoans, dinoflagellates, cnidarians, ctenophores, platyhelminths, tunicates and nematodes, but is absent from vertebrates, insects, plants and fungi [2]. Its phylogenetic distribution and rich molecular diversity suggests that it has evolved independently many times throughout eukaryote evolution [6–9], though an alternative scenario of early common origin with multiple losses may be possible [10].

One clear biological function of SL *trans*-splicing is the processing of polycistronic pre-mRNAs generated from eukaryotic operons [2]. In contrast to prokaryotes, where such transcripts can be translated immediately as they are transcribed, a key complication for eukaryotic operons is that nuclear polycistronic transcripts must be resolved into independent, 5’-capped monocistronic transcripts for translation in the cytoplasm [11]. The *trans*-splicing machinery coordinates cleavage of polycistronic pre-mRNA and provides the essential cap to the resulting un-capped monocistronic pre-mRNAs [12, 13]. This process is best characterised in the nematodes, largely, but not exclusively due to work on *C. elegans*, which possesses two types of SL [14]: SL1, which is added to mRNAs derived from the first gene in operons and monocistronic genes; and SL2, which is added to mRNAs arising from genes downstream in operons and thus specialises in resolving polycistronic pre-mRNAs [12–14].

The same SL2-type specialisation of some SLs for resolving downstream genes in operons exists in many other nematodes [15–20], but is not seen in other eukaryotic groups. For example, the platyhelminth *Schistosoma mansoni* and the tunicates *Ciona intestinalis* and *Oikopleura dioica* each possess only a single SL, which is used to resolve polycistronic mRNAs but is also added to monocistronic transcripts [21–23]. Similarly, the chaetognath *Spadella cephaloptera* and the cnidarian *Hydra vulgaris* splice a diverse set of SLs to both monocistronic and polycistronic transcripts [24, 25]. Remarkably, all protein-coding genes in trypanosomes are transcribed as polycistronic mRNAs and resolved using a single SL, making SL *trans*-splicing an obligatory component of gene expression [26]. In contrast, dinoflagellates use SL *trans*splicing for all nuclear mRNAs, but only a subset of genes are organised as operons [27, 28]. Although SL *trans*-splicing also occurs in many other organisms including rotifers, copepods, amphipods, ctenophores, cryptomonads and hexactinellid sponges, operons and polycistronic mRNAs have not been reported in these groups [7, 8, 29, 30].

All these examples illustrate a rich diversity in the SL *trans*-splicing machinery and its role in facilitating polycistronic gene expression and broader RNA processing. One barrier in dissecting the evolutionary history of these phenomena is the difficulty in systematically discovering novel SLs. Identifying the full SL repertoire of an organism would traditionally require laborious low-throughput cloning-based Sanger sequencing of the 5’ ends of mRNAs [e.g., 17, 31]. High-throughput RNA-Seq data is an attractive alternative resource that may often already exist for the focal organism. Novel SLs can, in principle, be identified from overrepresented 5’ tails extracted directly from RNA-Seq reads [32, 33]. Based on this idea, the SLFinder pipeline assembles putative SLs from overrepresented *k*-mers at transcript ends and uses these SLs as guides for searching potential SL RNA genes in genome assemblies [34]. SLFinder can detect known SLs in several eukaryotes but the assembled SLs are often incomplete or contain incorrect bases at the 3’ end [34]. Similarly, since SLFinder makes no assumptions about SL RNA structure apart from a GU splice donor site, the predicted SL RNA genes often contain pseudogenes [34].

An equally important barrier is the difficulty in quantifying SL *trans*-splicing events genome-wide and in establishing functional links between these events and operonic gene organisation. The 5’ ends of RNA-Seq reads contain, in principle, enough information to quantify SL *trans*-splicing events [29, 35], and genome-wide patterns of SL *trans*-splicing events have been exploited to predict novel operons from SL splicing ratios in the nematodes *Pristionchus pacificus* and *Trichinella spiralis* [19, 20]. The SL-QUANT pipeline automates quantification of SL *trans*-splicing events from RNA-Seq data for *C. elegans* and can be manually reconfigured to accept SL sequences and genomes from other organisms [36]. Similarly, the UTRme pipeline can identify and quantify 5’ UTRs associated with SL *trans*-splicing events by a single SL [37]. However, neither of these tools identify SLs specialised for resolving polycistronic mRNAs nor do they predict operons from SL *trans*-splicing events, and no other software exists to carry out these tasks.

Here we present two fully automated pipelines that address all these shortcomings of existing tools and present a unified and universal one-stop solution for systematically investigating SL *trans*-splicing and operonic gene organisation from RNA-Seq data in any eukaryotic organism. First, SLIDR is a more accurate, sensitive and specific alternative to SLFinder, implementing fully customisable and scalable *de novo* discovery of SLs and functional SL RNA genes. Second, SLOPPR is not only a more flexible and convenient alternative to SL-QUANT and UTRme for quantifying genome-wide SL *trans*-splicing events, but also uniquely provides algorithms for inferring SL subfunctionalisation and predicting operonic gene organisation. Both pipelines can process single-end or paired-end reads from multiple RNA-Seq libraries that may differ in strandedness, thus allowing for flexible high-throughput processing of large RNA-Seq or EST datasets from multiple sources.

## Implementation

### SLIDR: Spliced leader identification from RNA-Seq data

SLIDR is designed as an assembly tool that constructs initial putative SLs directly from 5’ read tails that remain unaligned (“soft-clipped”) after alignment to a genome or transcriptome reference [33]. Unlike other methods, SLIDR then implements several optional plausibility checks based on functional nucleotide motifs in the SL RNA molecule, i.e., splice donor and acceptor sites, *Sm* binding motifs and several stem loops [2, 38]. These features are expected to be present due to shared evolutionary ancestry of SL RNAs with the snRNAs involved in intron removal by *cis*-splicing [6, 39]. For each SL passing these filters, SLIDR reports read depth and annotates functionally plausible SL RNA genes and observed SL *trans*-splice acceptor sites in the reference.

RNA-Seq reads are first aligned to the genome or transcriptome reference using HISAT2 [40] or BOWTIE2 [41] in local alignment mode to enable soft-clipping at read ends. Since soft-clipped read tails must be long enough to capture full-length SLs (typically about 22 bp in nematodes), the alignment scoring functions are relaxed to allow for up to 35 bp tails in a 100 bp read by default (HISAT2: `--score-min L,5,−0.4 --sp 1,0 --mp 3,1`; BOWTIE2: `--score-min L,5,0.6 --ma 1 --mp 3,1`). A scaling factor can be supplied by the user to restrict or expand the upper limit to accommodate more extreme SL lengths, for example 16 bp in *Ciona intestinalis* [42] or 46 bp in *Hydra vulgaris* [25]. Tails from the read end corresponding to the 5’ end of the transcript (inferred from library strandedness) are extracted using SAMTOOLS [43].

The extracted tails are then dereplicated, 3’-aligned and clustered at 100 % sequence identity using VSEARCH [44]. Each read cluster thus represents a single putative SL, comprising a collection of 3’-identical read tails that differ in 5’ extent (Fig. 1). VSEARCH provides two clustering modes, assigning each tail to the cluster with either the longest (distance-based greedy clustering; DGC) or most abundant centroid (abundance-based greedy clustering; AGC) that matches at 100 % identity [44, 45]. In our experience, the default DGC is usually superior, but we have observed cases where AGC recovers magnitudes more reads because DGC clustered large numbers of very short tails with a long, noisy centroid that failed to align to the genome. SLIDR provides the option to use AGC if required, which is only advisable when the final SLs have suspiciously low read coverage.

**Fig. 1:**
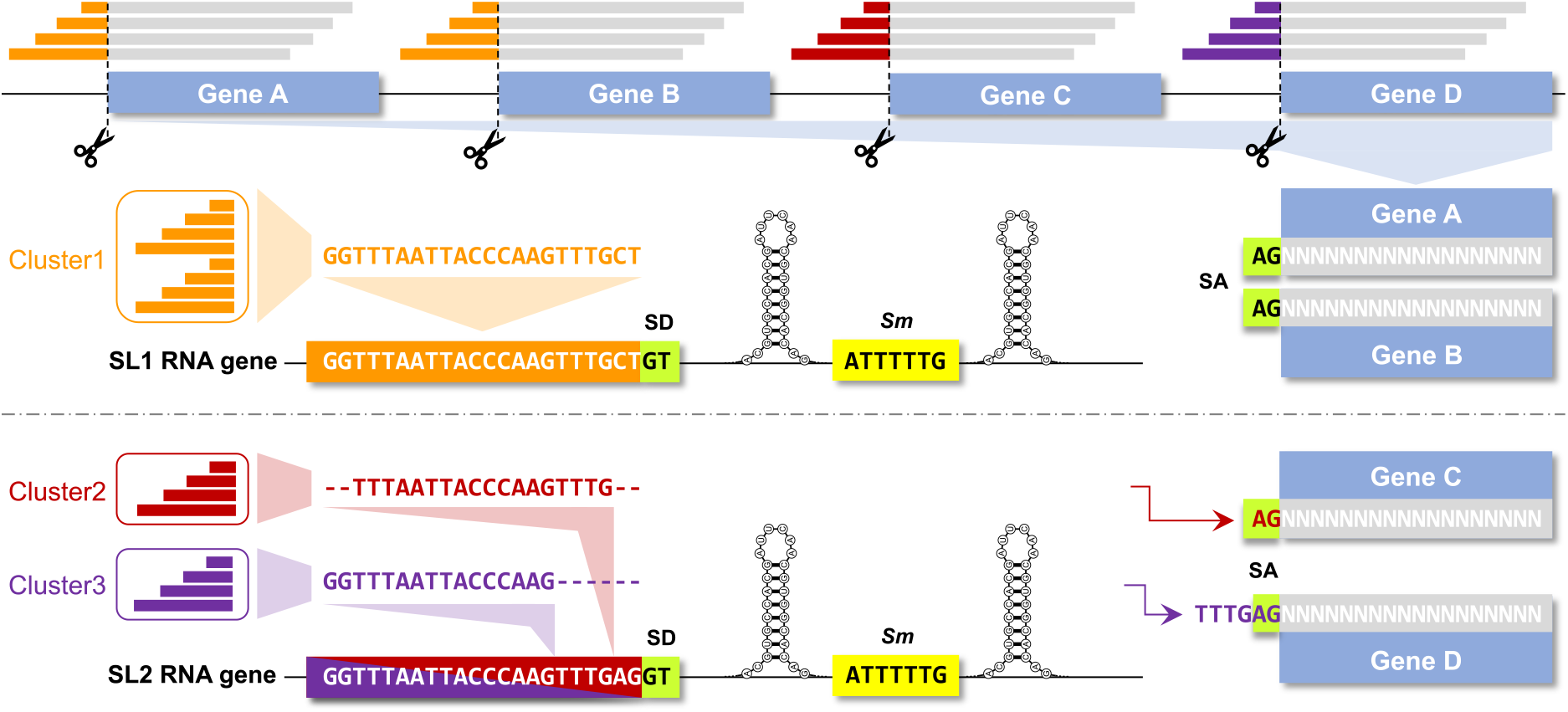
Schematic representation of the SLIDR pipeline (Spliced leader identification from RNA-Seq). Local alignments of reads (grey) to a genomic reference (illustrated by four genes A-D) allow for 5’ spliced leader (SL) tails to be soft-clipped and extracted (coloured read portions). Clustering of 3’ aligned read tails from all genes at 100% sequence similarity produces unique consensus SL candidates (cluster centroids), which are required to align to the genomic reference to identify candidate SL RNA genes (illustrated by SL1 and SL2 genes). In SL RNA genes, a splice donor site (SD; for example GT) is expected immediately downstream of the genomic alignment, followed by an *Sm* binding site (for example 5’-ATTTTTG-3’) bookended by inverted repeats capable of forming stem loops in the RNA transcript. Conversely, the spliced gene requires a splice acceptor site (SA; for example AG) immediately upstream of the 5’ read alignment location in the genomic reference. In this illustration, the example SL1 is fully reconstructed from a single read-tail cluster (cluster 1) with GT and AG splice sites in the expected locations (genes A and B). In contrast, the example SL2 highlights how read tails may be 3’-truncated due to overlap with the splice acceptor site (genes C and D) and the upstream *trans*-splice acceptor site sequence at some genes (gene D). These missing nucleotides can be filled in from the *trans*-splice acceptor site region guided by the distance between the 3’ tail alignment location and the splice donor site (GT). Note that although cluster 2 is also 5’ truncated due to insufficient coverage at gene C, consensus calling with cluster 3 allowed for reconstructing the full SL2 RNA gene.

The cluster centroids are then used to identify candidate SL RNA genes in the genome or transcriptome reference. The centroids are aligned to the reference using BLASTN [46] with 100% sequence identity and a relaxed customisable E-value of 1 to accommodate short queries. Matches are required to contain the full 3’ end of the centroid but may be 5’ truncated to allow for 5’ noise in RNA-Seq reads. For each match, the full putative SL RNA gene sequence (of customisable length) is extracted from the reference using BEDTOOLS [47]. This sequence is then inspected for customisable splice-donor (default: GT) and *Sm* binding (default: AT{4,6}G) sites, and secondary structure stem loops are predicted using RNAFOLD from the ViennaRNA package [48]. Default criteria expect the *Sm* motif 20-60 bp downstream of the splice donor site [49]. In the reference sequence immediately upstream of the aligned portion of each RNA-Seq read, a splice acceptor site (default: AG) is required, corresponding to the SL *trans*-splice acceptor site of the gene (Fig. 1).

The locations of splice donor and *trans*-splice acceptor sites may not be as expected if the 3’ end of the SL and the 3’ end of the *trans*-splice acceptor site happen to be identical. In these cases, the RNA-Seq read alignment overextends in 5’ direction into the *trans*-splice acceptor site and thus 3’-truncates the soft-clipped SL read tail (Fig. 1). These missing 3’ nucleotides can be reconstructed from surplus nucleotides located between the 3’ end of the centroid BLASTN match and the splice donor site, and must be identical to those surplus nucleotides located between the 5’ read alignment location and the splice acceptor site (Fig. 1). Following reconstruction of the 3’ end where necessary, all tail cluster centroids are subjected to another round of 3’ alignment and clustering at 100% sequence identity in VSEARCH before final SL consensus construction is carried out in R [50]. Final SLs must be supported by at least two reads and must be spliced to at least two genes that are not located in the immediate vicinity (1 kbp distance) of the SL RNA gene [32].

### SLOPPR: Spliced leader-informed operon prediction from RNA-Seq data

SLOPPR is designed as a genome annotation tool that predicts operons from genome-wide distributions of SL *trans*-splicing events at pre-annotated genes. RNA-Seq reads that contain evidence of 5’ SLs are identified using a sequence-matching approach equivalent to the “sensitive” mode of SL-QUANT [36]. The operon prediction algorithm is built upon the SL1/SL2-type functional specialisation of SLs observed in many nematodes, but is fully customisable to accommodate other relationships between SLs and operonic genes, even when SL specialisation is absent. Unlike previous approaches that have defined operons in various organisms primarily via short intercistronic distances [18, 19, 22, 51], SLOPPR defines operons principally via SL *trans*-splicing patterns and only optionally takes intercistronic distance into account. SLOPPR can also identify and correct gene annotations where operonic genes are incorrectly annotated as a fused single gene [20, 52], paving the way for *trans*-splicing-aware genome (re-)annotation.

RNA-Seq reads containing SLs are identified using a three-step strategy equivalent to SL-QUANT [36]. Since such reads cannot align end-to-end to the genome because of the *trans*-spliced 5’ SL tail, all reads are first aligned end-to-end to the genome reference using HISAT2 [40] and unmapped reads are retained as candidates. If paired-end reads are used, the read corresponding to the 3’ end of the transcript (inferred from library strandedness) must be aligned, and the read corresponding to the 5’ end of the transcript must be unaligned [36]. The 5’ ends of the unaligned candidate reads are then screened for overlap with the 3’ portion of any number of supplied SL sequences using CUTADAPT [53]. Finally, those reads that align to the genome end-to-end after the SL tail has been trimmed are quantified against exons and summarised at the gene level using FEATURECOUNTS from the SUBREAD package [54]. Likewise, background expression levels of all genes are obtained from the original end-to-end read alignments and from candidate reads without SL evidence. This screening strategy is carried out for each RNA-Seq library independently, thus allowing for comparisons among biological replicates during analysis (Fig. 2A).

**Fig. 2:**
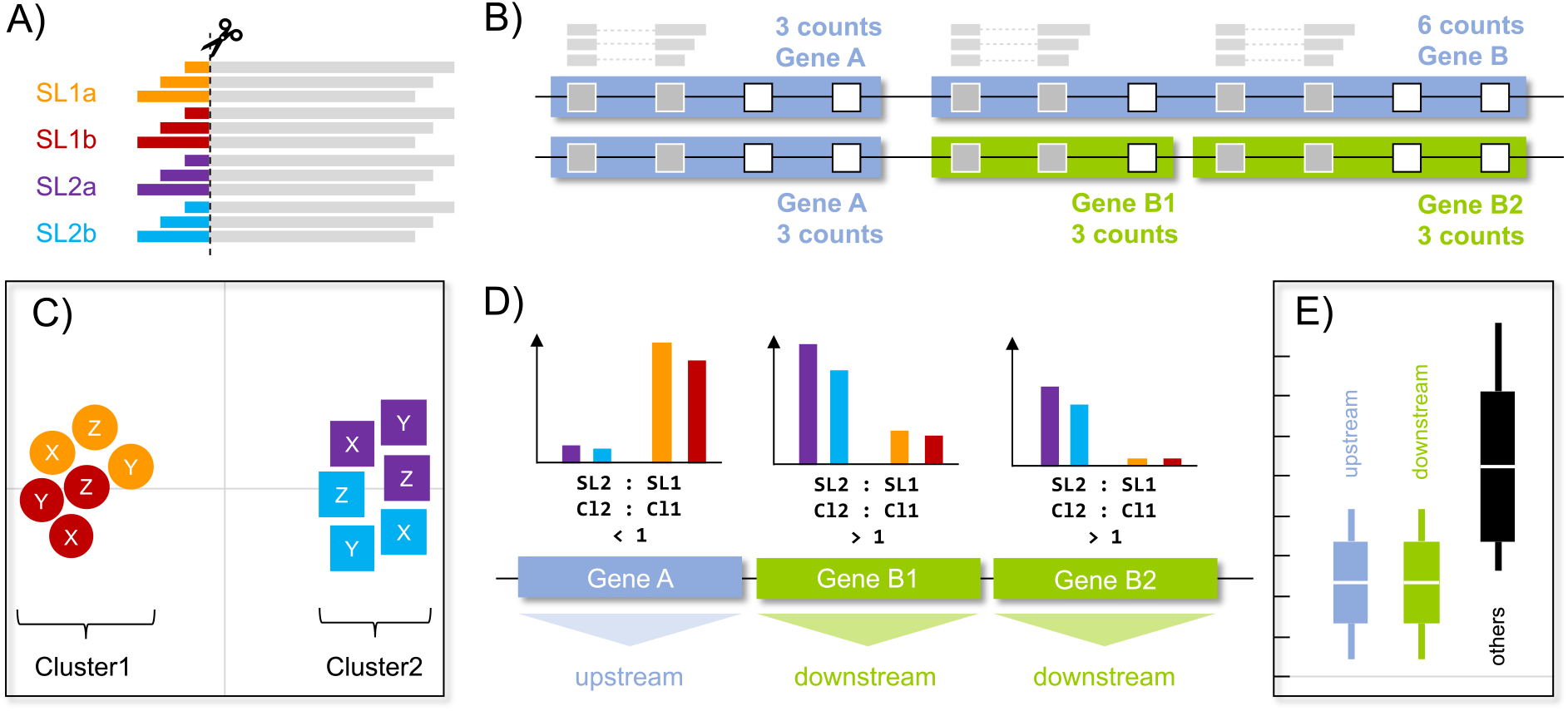
Schematic representation of the SLOPPR pipeline (Spliced leader-informed operon prediction from RNA-Seq). A) Spliced leader tails (example: SL1a, SL1b, SL2a and SL2b) are identified and trimmed from the 5’ end of reads that correspond to the 5’ end of transcripts. B) Trimmed reads are aligned to the genome, quantified against exons (squares; grey: covered; white: not covered) and counts are summarised by gene (example: two genes A and B). Incorrect gene annotations (fused operonic genes) can optionally be identified and corrected via SL reads at internal exons (example: Gene B is split into B1 and B2). C) SL read sets from multiple libraries (example: X, Y and Z) are ordinated via PCA on genome-wide read counts and grouped into two clusters (K-means clustering) expected to correspond to SL1 (circles) and SL2-type (squares) subfunctionalisation. D) SL2:SL1 read ratios are computed between pre-defined SL groups (SL1, SL2) or inferred clusters (Cl1, Cl2). Operons are predicted via tracts of genes receiving SL2 bias (downstream operonic genes) plus an optional upstream gene receiving either an SL1 bias or no SLs at all. E) Intercistronic distances among predicted operons are expected to be reduced compared to intergenic distances among non-operonic genes (others). Operon predictions can optionally be filtered by intercistronic distance using a user-supplied or inferred optimal cutoff.

The nature of the SL *trans*-splicing process means that SLs must only be present at the first exon of a gene, i.e. the 5’ end. We can thus identify problematic gene annotations that consist of fused operonic genes via internal exons that receive SL reads [20, 52]. SLOPPR implements an optional gene-correction algorithm that splits gene annotations at exons with distinct SL peaks compared to neighbouring exons (Fig. 2B). To obtain exon-based SL counts, genome annotations are converted to GTF using GFFREAD from CUFFLINKS [55], unique exons are extracted using BEDTOOLS [47] and SL reads are quantified with FEATURECOUNTS at the exon level instead of gene level. The peak-finding algorithm is designed to correctly handle reads that may span multiple exons (Fig. 2B).

The SL read counts obtained from FEATURECOUNTS are normalised for library size using CPM (counts-per-million) based on the background gene counts [56]. The normalised SL read-count matrix is then subjected to generalized principal component analysis (PCA) and hierarchical clustering designed for sparse count matrices [57], treating SL read sets as samples and genes as variables. This summary of genome-wide distributions of SL *trans*-splicing events allows for identifying the distinct *trans*-splicing patterns of SL2-type SLs expected from their specialisation to resolve downstream operonic genes. If SL2-type SLs are not known, K-means clustering and linear discriminant analysis are used to assign SLs to one of two synthetic clusters assumed to correspond to SL1-type and SL2-type SLs (Fig. 2C). Visual inspection of the clustering results allows the user to determine consistency across biological replicates (if available) and to ascertain functional groups of SLs.

Based on the SL clustering results and pre-defined SL1/SL2-type groups (if known), the SL2:SL1 CPM ratio is computed and summarised across all genes that receive both SL types. The operon prediction algorithm is based on finding uninterrupted runs of adjacent genes with SL2-bias, which are designated as downstream operonic genes (Fig. 2D). By default, no SL1-type reads are allowed at downstream genes (i.e., SL2:SL1 = infinity), but a more relaxed SL2:SL1 ratio cutoff can be provided. The optimal cutoff is species-specific and could be identified empirically from inspecting the distribution of SL2:SL1 read ratios or from observed read ratios at known operonic genes [20]. Each tract of SL2-biased operonic genes then receives an upstream operonic gene that shows SL1-type bias or absence of SL *trans*-splicing (Fig. 2D). Alternatively, upstream operonic genes can be required to have the same SL2-type bias as downstream genes.

Finally, intercistronic distances among the predicted operonic genes are computed and compared to genome-wide intergenic distances to diagnose tight physical clustering of operonic genes (Fig. 2E). These distances are obtained from the boundaries of consecutive “gene” GFF annotation entries, so their accuracy depends entirely on the provided genome annotations, which should ideally define gene boundaries by poly(A) and *trans*-splice acceptor sites. If desired, operon prediction can take intercistronic distances into account, either via a user-supplied distance cutoff or via an automatic K-means clustering method that splits the genome-wide distribution of intercistronic distances into two groups, corresponding to tight gene clusters (potential operons) and non-operonic genes. In consequence, by manually specifying SL1/SL2-type SLs, SL2:SL1 ratio cutoff, upstream gene SL2-type bias and intercistronic distance cutoff, a large gamut of relationships between SLs and operonic genes can be explored, even in situations where no subfunctionalisation of SLs for operon resolution exists, for example in kinetoplastids or tunicates [21–23].

## Results and Discussion

### SLIDR validation and comparison with SLFinder

We first compared the accuracy of SLIDR against that of SLFinder [34], focusing on *C. elegans, Ciona intestinalis, Hydra vulgaris* and *Schistosoma mansoni*, and using the same reference genomes and RNA-Seq datasets that were used in benchmarking SLFinder [34]. All RNA-Seq datasets were quality-filtered with TRIM_GALORE 0.6.4 [58], trimming Illumina adapters (5’-AGATCGGAAGAGC-3’) and poor-quality bases (phred 20) from 3’ read ends. All SLIDR runs required the presence of the canonical GT/AG splice donor/acceptor sites. For each organism, we examined how many of the expected SLs and SL RNA genes were detected (sensitivity), whether the SL sequence was accurate and complete, how many genes were SL *trans*-spliced (SL *trans*-splicing rate) and whether any novel SL candidates were identified.

The results demonstrate that SLIDR detects expected SLs with greater sensitivity and accuracy than SLFinder and performs well across eukaryotes with SL repertoires and SL RNA structures that are divergent from *C. elegans* (Table 1). SLIDR assembled accurate and complete SLs, whereas those assembled by SLFinder contained incorrect nucleotides at the 3’ end and were 5’ truncated in one case (Table 1). SLIDR identified smaller sets of SL RNA genes than SLFinder because SLIDR only reports potentially functional loci given the RNA-Seq data, whereas SLFinder is designed to annotate all possible gene loci given initial SLs (“hooks”) assembled from transcript ends in *de novo* transcriptome assemblies [34]. Both approaches are clearly complementary, though we find that SLIDR’s focus on evidence from RNA-Seq reads and functional plausibility is more effective in detecting accurate SLs and SL RNA genes (Table 1).

**Table 1:**
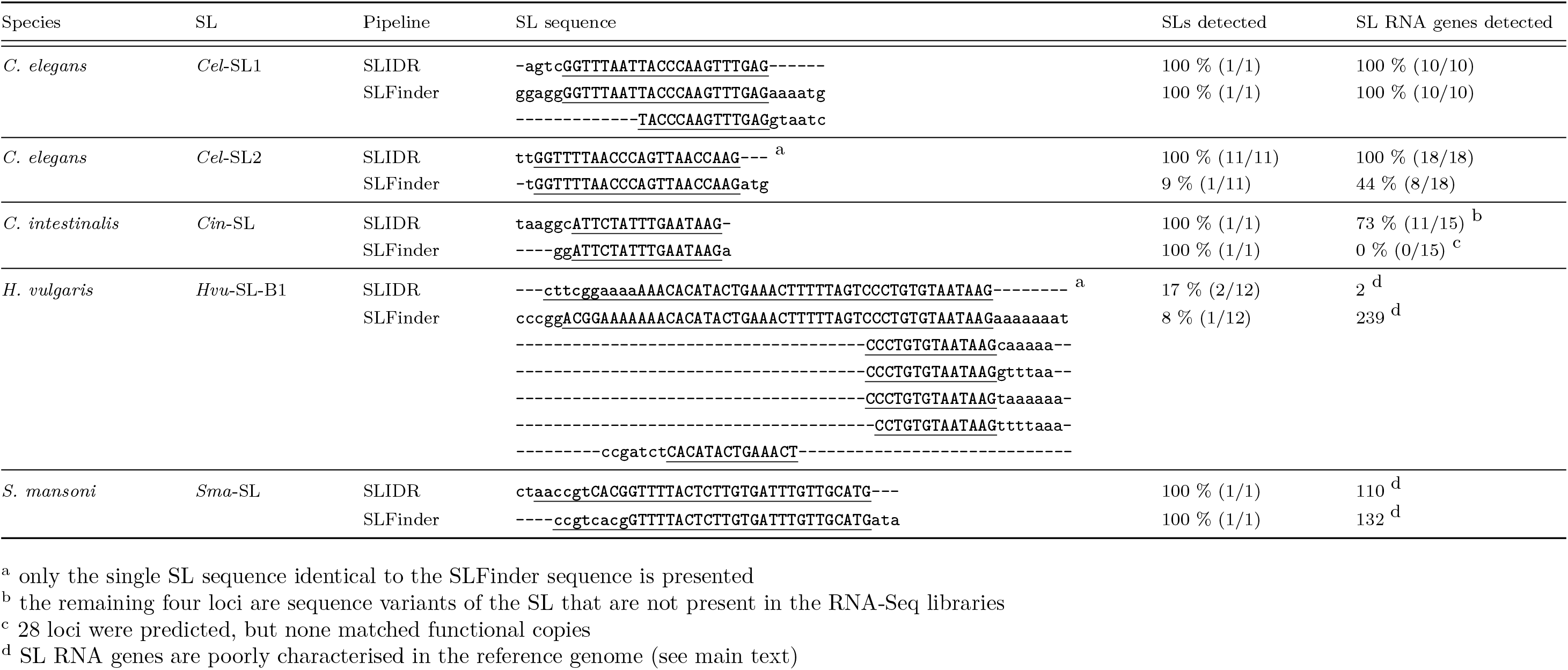
Comparison of SLs and SL RNA genes detected by SLIDR and SLFinder in four eukaryotes (*Caenorhabditis elegans, Ciona intestinalis, Hydra vulgaris* and *Schistosoma mansoni*). Datasets are detailed in Supplemental Table S1 and [34]. For each expected SL, the sequences assembled from the RNA-Seq data are presented as alignments, underlining the expected sequence. Lower-case 5’ bases in the SLIDR sequences denote bases supported by fewer than 50 % of reads. Numbers of detected out of expected SLs and SL RNA genes are detailed where available.

#### Caenorhabditis elegans

*C. elegans* possesses the best understood repertoire of nematode SLs, comprising two types, SL1 and SL2, both of which are encoded by multi-copy gene families with well-described SL RNA structures and *Sm* binding motifs [12–14, 56]. SL *trans*-splicing affects up to 84 % of genes [35, 56], which makes *C. elegans* an appropriate benchmark organism for SL detection pipelines. We ran SLIDR with parameter `-S’.{40,55}AC?T{4,6}G’` to reflect the *Sm* binding site motif [38, 49] and compared the identified SL RNA genes with the reference gene annotations using BEDTOOLS INTERSECT 2.28.0 [47].

SLIDR assembled 141,019 reads into the correct SL1 sequence, and detected ten SL RNA genes, corresponding to all ten functional *sls-1* genes (*sls-1.1* and *sls-1.5* are 5’-truncated pseudogenes that are incorrectly annotated in the assembly). SLIDR also identified all eleven SL2 sequence variants from as few as 28 reads and detected 18 SL RNA genes, which correspond to all 18 functional *sls-2* genes and correctly omitted the *sls-2.19* pseudogene (Supplemental Table S1). These results illustrate 100 % sensitivity for both SLs and SL RNA genes. A total of 11,712 genes received SLs, yielding a 58 % SL *trans*-splicing rate (Supplemental Table S1). Interestingly, SLIDR also reported 25 additional sequences that behaved like an SL, of which 13 were derived from annotated genes with diverse functions. These results may be evidence of genic *trans*-splicing events [2] or may be chimeric artefacts since most of them were supported by only two reads (Supplemental Table S1).

In comparison, SLFinder also correctly identified the ten functional *sls-1* genes but rarely identified splice donor sites and also reported seven pseudogenes [34]. Strikingly, SLFinder detected only five out of eleven SL2 sequence variants and only eight out of 18 *sls-2* genes [34]. This might be because only three initial SL sequence “hooks” were assembled from the transcriptomes, and these matched only a single SL2 sequence variant [34]. Although all three hooks matched SL sequences (100% specificity), the hook sequences were inaccurate because they were either 5’ truncated or contained up to six unspecific nucleotides at the 3’ end (Table 1). While these results are certainly sufficient for discerning correct SLs and SL RNA genes with manual curation [34], SLIDR yields accurate results even from few reads.

#### Ciona intestinalis

The tunicate *C. intestinalis* possesses a single 16 bp spliced leader 5’-ATTCTATTTGAATAAG-3’ that is spliced to at least 58% of expressed genes [22, 42, 59]. The SL RNA is very short (46 bp), contains the *Sm*-binding motif 5’-AGCUUUGG-3’ [60] and is encoded by a highly repetitive gene family comprising at least 670 copies, though the reference genome contains at most 15 of them due to assembly constraints [61]. We ran SLIDR with the parameters `-x 0.6 −e 5 −O 5 −R 30 −S ’.{2,25}AGCTTTGG`’ to enforce shorter soft-clipping (maximum 24 bp given 100 bp reads), a BLAST e-value cut-off of 5 (to allow short matches of c. 11 bp), maximum 5 bp overlap with the *trans*-splice acceptor site, 30 bp RNA length excluding the SL, and the *Sm*-like motif located up to 25 bp downstream of the GT splice donor site.

SLIDR identified the expected SL from only 95 reads (spliced to 93 genes) despite very high genome alignment rates of 93-95 %. The SL sequence contained extra 5’ nucleotides supported by the minority of reads (5’-taaggcATTCTATTTGAATAAG-3’). All but one of the eleven SL RNA genes identified by SLIDR were on chromosome NC_020175.2 (one was on NC_020166.2), and all were part of a 264 bp repeat unit that contains functional SL copies in the genome [61]. In comparison, SLFinder assembled a single hook after relaxing assembly parameters and hook filters, which matched the correct SL sequence with only 1 unspecific nucleotide at the 3’ end (Table 1). This hook yielded two distinct gene variants comprising 28 putative SL RNA genes, of which 21 had a splice donor site [34]. None of these genes were located within the 264 bp repeat unit and may therefore be pseudogenes, which are rife in this organism [61].

To explore the cause of the poor read coverage in the SLIDR analysis, we first re-ran SLIDR removing the filter for the *Sm* motif, which identified the same SL at identical coverage but yielded >50 additional SL RNA genes (likely pseudogenes) (Supplemental Table S1). Similarly, two additional runs using libraries from two other bioprojects yielded similar coverage (29-150 reads) and no more than 91 SL *trans*-spliced genes (Supplemental Table S1). However, using 13 libraries from a final bioproject yielded 70,745 reads spliced to 7,160 genes and originating from the same eleven SL RNA genes as above. Additionally, two novel SL variants were detected from 13-378 reads spliced to 15-281 genes and originating from three novel SL RNA genes, totalling 14 out of 15 expected genes [61] (Supplemental Table S1). Assuming 15,254 genes in the genome [22], these results yield a SL *trans*-splicing rate of 47 %, which is much closer to the expected 58 % [59]. This substantial variability of SL *trans*-splicing rates between biosamples may be due to variability among life stages, tissues or RNA-Seq library preparation methods.

Finally, SLIDR also discovered potentially novel SLs that resemble nematode SLs instead of the canonical *C. intestinalis* SL. After removing the *Sm* motif filter during the analysis of the original libraries above we noticed a novel putative SL supported by 3,612 reads (Supplemental Table S1). We re-ran SLIDR with the default parameters designed for nematode SLs and confirmed a novel 21 bp SL (5’-CCGTTAAGTGTCTTGCCCAAG-3’) defined by 3,621 reads but spliced to only 7 genes (Supplemental Table S1). That same novel SL was also detected among the 13 libraries of bioproject PRJNA376667, though at much lower coverage of only 24 reads (Supplemental Table S1). It was beyond the scope of this study to fully resolve and describe this novel SLs, but these preliminary results do highlight that SLIDR is more sensitive than SLFinder, which found no evidence of this SL in the original libraries [34].

#### Hydra vulgaris

The cnidarian *H. vulgaris* possesses two types of SLs that are added to at least one third of all genes: the first type (SL-A) is 24 bp long and is part of an 80 bp SL RNA [62], whereas the second type is longer (46 bp SL, 107 bp SL RNA) and comprises a total of eleven SL variants across six SLs (SL-B to SL-G) [25]. The *Sm* binding sites differ between SL-A (5’-GAUUUUCGG-3’) and all other SLs (5’-AAUUUUGA-3’ or 5’-AAUUUUCG-3’) [62]. We ran SLIDR with the parameters `-x 1.5 −R 60 −S ’.{l0,35} [AG]ATTTT [CG][AG]`’, which cover both *Sm* binding site motifs and should allow for detecting both the short and long SLs.

SLIDR detected the full SL-B1 sequence from 865,003 reads spliced to 18,768 genes and identified two SL RNA genes. SLIDR also detected two 5’ truncated versions of SL-D (encoded by two SL RNA genes) and two potential novel SL variants at much lower coverage (Supplemental Table S1). SLFinder assembled six hooks, all of which matched SL-B1 but were either considerably 5’ truncated or contained up to eight unspecific nucleotides at the 3’ end (Table 1). These hooks yielded 239 putative SL RNA genes, which included the two SL-B1 genes identified by SLIDR, but comprised mostly 5’ truncated genes that were missing splice donor sites and are thus probably pseudogenes [34].

These results highlight that SLIDR is more sensitive and specific than SLFinder but still missed ten out of twelve SLs and did not recover the full 5’ extent of SL-D. Since these SLs are exceptionally long, reads longer than 100 bp may be required to detect full-length SLs. We thus tested SLIDR with 2×150 bp reads from a different bioproject and detected full-length SL-B1, SL-D and SL-E based on 6,773–294,822 reads spliced to 1,885–13,325 genes, and two novel SL-B-type variants at much lower coverage (Supplemental Table S1). Contrary to previous estimates that only about 33 % of about 20,000 protein-coding genes are SL *trans*-spliced [25], both SLIDR runs suggest that at least 67-94 % (13,325-18,768) genes may be SL *trans*-spliced.

#### Schistosoma mansoni

The platyhelminth *S. mansoni* possesses a single, relatively long (36bp) SL with an unusually long *Sm* binding site (5’-AGUUUUCUUUGG-3’) and a total SL RNA length of 90 bp [63]. The transcripts from at least 46 % of genes undergo *trans*-splicing by this SL [23]. SLIDR was run with parameters `-x 1.25 −R 55 −S ’.{10,30}AGTTTTCTTTGG`’ to allow for detecting this large SL and the *Sm* binding site.

SLIDR detected the complete SL with two extra 5’ nucleotides (5’-ctaaccgtcacgGTTTTACTCTTGTGATTTGTTGCATG-3’) supported by 41,243 reads, and two 5’-truncated SL variants supported by only 3 reads each (Supplemental Table S1). The canonical SL was encoded by 110 SL RNA genes, of which all except one were tightly clustered on chromosome SM_V7_6. This is consistent with the presence of a repeat unit comprising as many as 200 SL RNA gene copies [63], though the genome annotations contain only five curated copies [23]. A total of 3,746 genes (30 %) were SL *trans*-spliced, which exceeds the 2,459 genes previously identified from large-scale RNA-Seq data (250 million reads; [23]), but falls short of the expected 46 % SL *trans*-splicing rate [23].

SLFinder assembled a single hook that missed the two 5’ A nucleotides and included three unspecific 3’ nucleotides (Table 1). All 132 detected SL RNA genes were considerably 5’ truncated but all except nine had a clear splice donor site [34]. SLFinder detected the same 110 genes as SLIDR, notwithstanding incorrect 5’ truncation caused by the trimming algorithm [34]. Interestingly, SLFinder detected a genomic locus where the terminal ATG nucleotides of the SL sequence were replaced by ACG [34]; SLIDR did not detect this variant because it was not informed by evidence from the RNA-Seq data. Overall, both tools yielded comparable results, but SLIDR omitted likely pseudogenes and detected the full 5’ extent of the SL sequence and the SL RNA genes.

### SLIDR performance in nematodes

Having validated SLIDR in *C. elegans* and three other eukaryotes, we then examined SLIDR’s performance (using the same criteria as above) in a range of other nematodes with well-characterised SL and SL RNA repertoires: *Caenorhabditis briggsae, Pristionchus pacificus, Meloidogyne hapla, Trichinella spiralis* and *Trichuris muris*. We also tested how SLIDR performs with a transcriptome reference instead of a genome. In this situation, SLIDR cannot confirm splice acceptor sites because these are not expected to be present in a transcriptome; however, if the transcriptome contains SL RNAs it will still be possible to find splice donor and *Sm* binding sites. We demonstrate this in *Prionchulus punctatus* using a draft *de novo* transcriptome assembly and compare the results against *C. elegans* using a well-curated transcriptome.

As above, all RNA-Seq data underwent quality-trimming and SLIDR required the default GT/AG splice donor/acceptor sites. The results demonstrate that SLIDR detects all known SLs in all cases and often discovers novel SL variants (Table 2). We also illustrate that the transcriptome mode of SLIDR works well with curated transcriptomes, and can, in principle, identify SLs even from draft transcriptomes assembled *de novo* from the focal RNA-Seq data itself (Table 2). While the genome-mode is highly preferred, the transcriptome mode allows to gain initial insight into SL repertoire even for organisms with poor or absent genomic resources.

**Table 2:**
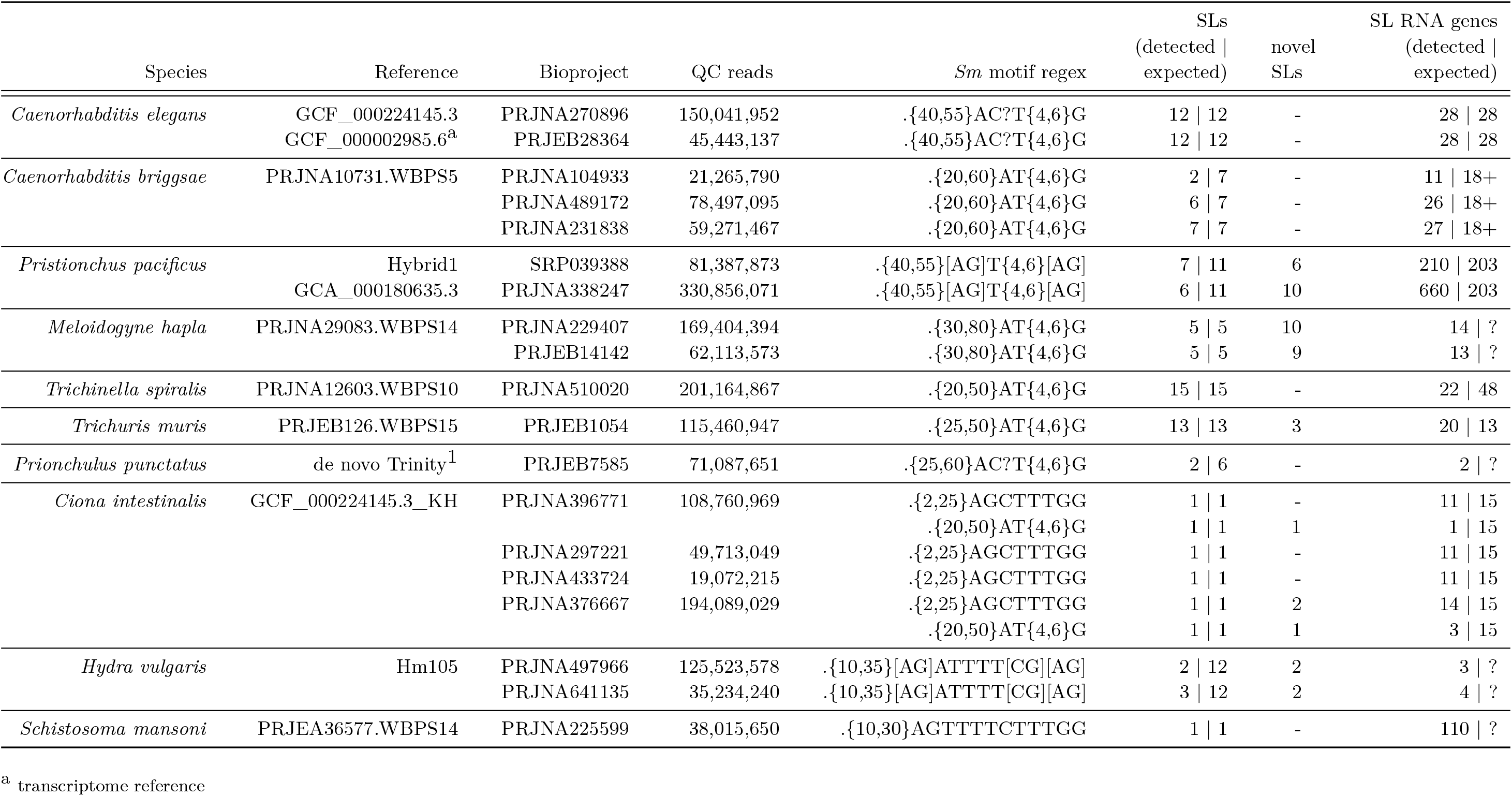
Spliced leader identification (SLIDR) results in seven nematodes and three other eukaryotes. Identifiers for reference genomes/transcriptomes and RNA-Seq libraries are presented alongside numbers of quality-trimmed reads (QC), the *Sm* motif regular expression notation used to filter SLs, numbers of expected SLs detected, numbers of novel SLs identified and numbers of expected SL RNA genes detected. Question marks and pluses represent unknown or poorly characterised SL RNA gene numbers.

#### Caenorhabditis briggsae

*C. briggsae* is a close relative of *C. elegans* that possesses a similar SL repertoire and shows considerable synteny of operons [16, 18]. *Cbr*-SL1 is encoded by a repetitive gene cluster (about 65 copies; 64) linked to 5S rRNA genes [65], whereas SL2-type SLs are encoded by 18 genes and represent six distinct SL variants (*Cbr*-SL2, *Cbr*-SL3, *Cbr*-SL4, *Cbr*-SL10, *Cbr*-SL13, *Cbr*-SL14), of which four are shared with *C. elegans* [16]. Only 37 % of genes are SL *trans*-spliced [18], compared to 70–84 % in *C. elegans* [35, 56], though this is likely simply a reflection of differential transcriptome read depth used in these studies.

We first used the same 2×42 bp RNA-Seq data that were used originally to identify genome-wide SL *trans*-splicing events [18]. These data are particularly difficult to analyse because the short reads are likely to impede identification of the full-length SL. To maximise SL detection, SLIDR was run with parameters `-S ’.{20,60}AT{4,6}G’ −x 2`, which would allow a tail of at most 28 bp and leave at least 14 bp for read alignment. Irrespective, SLIDR only detected 5’ truncated versions of *Cbr*-SL1 (2,377 reads spliced to 743 genes) and *Cbr*-SL3 (17 reads spliced to 17 genes) (Supplemental Table S1).

We then tried a stranded 2×50 bp library with more than three times as many reads as above, and detected full-length *Cbr*-SL1 (110,401 reads spliced to 7,703 genes) and five out of six full-length SL2-type SLs (*Cbr*-SL13 was absent) supported by up to 20,317 reads and spliced to up to 1,968 genes (Supplemental Table S1). Finally, using data from five unstranded 2×76 bp libraries, SLIDR recovered *Cbr*-SL1 (149,882 reads spliced to 8,054 genes) and all six SL2-type SLs (6–72 reads spliced to 6–42 genes) (Supplemental Table S1).

Across these three sets of libraries, SLIDR identified four SL RNA genes for *Cbr*-SL1 and 23 instead of 18 genes for SL2-type SLs [18, 65]. These results illustrate that SLIDR can detect SLs and SL RNAs even from very short reads in an organism where relatively few genes are SL *trans*-spliced (Table 2).

#### Pristionchus pacificus

*P. pacificus* possesses seven SL1-type (*Ppa*-SL1) and four SL2-type (*Ppa*-SL2) SLs, which are encoded by 187 and 16 genes respectively [19]. Based on *Ppa*-SL1a-and *Ppa*-SL2a-enriched RNA-Seq, about 90 % of 23,693 expressed genes are SL *trans*-spliced [19]. We ran SLIDR independently on those same three libraries (SL1-enriched, SL2-enriched and non-enriched control) [19] to examine how SL-enriched RNA-Seq affects SLIDR’s performance compared to non-enriched data. For all runs, we specified the parameter `-S ’.{40,55}[AG]T{4,6}[AG]`’ to capture both SL1 and SL2 *Sm* binding motifs [15].

The non-enriched library yielded substantial evidence for *Ppa*-SL1a (97,630 reads spliced to 4,328 genes) and little evidence of *Ppa*-SL2a (40 reads spliced to 16 genes). The SL1-enriched library yielded *Ppa*-SL1a at double the coverage (190,892 reads spliced to 6,443 genes) and *Ppa*-SL2a/m at low coverage (at most 140 reads), consistent with SL1-enrichment. However, the SL2-enriched library contained predominantly *Ppa*-SL1a (52,697 reads spliced to 3,110 genes) despite increased coverage of *Ppa*-SL2a (4,656 spliced to 604 genes), suggesting that the SL2-enrichment may not have been as effective as suggested by qPCR control experiments [19]. Across the three libraries, SLIDR detected three out of seven *Ppa*-SL1 variants, all four *Ppa*-SL2 variants and six novel SL variants (Supplemental Table S1). Overall, 188 SL1 RNA genes and at least 22 SL2 RNA genes were detected (Table 2), which is a slight increase compared to previous reports [19], but the estimated SL *trans*-splicing rate of 38 % falls way short of the anticipated 90 % [19].

To explore how SLIDR performs with different data, we then used six unstranded 2×150 bp libraries from bioproject PRJNA338247 and a more recent, better resolved genome assembly (GCA_000180635.3). SLIDR detected all but one of the same known SL1 and SL2 variants as before, with similar coverage and numbers of SL *trans*-spliced genes (Supplemental Table S1). Due to the superior genome assembly, SLIDR detected at least 619 SL1 RNA genes and at least 41 SL2 RNA genes; many more than previously reported (Table 2). SLIDR also detected at least ten novel SL variants, with large numbers of plausible additional SL variants at low read depths among large amounts of noise (Supplemental Table S1). This suggests that more extensive RNA-Seq datasets are required to fully resolve the landscape of SLs and SL RNA genes in this organism.

These results highlight a rich diversity of known and previously unreported SL1-type and SL2-type SLs beyond the canonical *Ppa*-SL1a and *Ppa*-SL2a variants [16, 19]. The striking discrepancy in observed versus expected SL *trans*-splicing rate can, in part, be explained by SLIDR’s reliance on a small fraction of RNA-Seq reads that contain sufficient evidence of an SL tail at their 5’ end. Any underrepresentation of SL tails in the original libraries [19] may be due to 5’ bias caused by obsolete library preparation chemistry [66]. Conversely, the SLIDR results may also point to issues with the SL enrichment underpinning the original SL *trans*-splicing rate estimates [19]. If the enriched libraries were contaminated with non-*trans*-spliced transcripts, the SL *trans*-splicing rate would be overestimated. Since this is impossible to test bioinformatically with the available data, more work is required to verify the extent of SL *trans*-splicing in this organism.

#### Meloidogyne hapla

The plant-root knot nematode *M. hapla* possesses the canonical *C. elegans* SL1 and four additional variants, all of which are *trans*-spliced to a minority of only 10 % of 14,420 genes [32]. We obtained the same 32 libraries that were used to discover these SLs [32]. These data are particularly difficult to analyse because they are 75 bp single-end, unstranded and originate from mixed-culture RNA samples containing primarily material from the host plant *Medicago truncatula*. Since reads from unstranded single-end libraries originate from the 5’ end of the transcript only 50% of the time, usable coverage is effectively halved. SLIDR was run with parameters `-S ’ .{30,80}AT{4,6}G’ −R 90` to allow for larger variation in *Sm* binding motif location and a longer SL RNA.

SLIDR detected all five known SLs and discovered at least ten novel SLs, suggesting that the SL repertoire in this organism is much larger than previously identified (Supplemental Table S1). The SLs were supported by up to 28,544 reads and were spliced to up to 5,742 genes (Supplemental Table S1). This indicates an SL *trans*-splicing rate of at least 40 %, which suggests that previous reports of 10 % is an underestimate [32]. We confirmed this with different RNA-Seq data (100 bp single-end), whereby SLIDR detected the same known and novel SLs with comparable coverage and up to 6,928 *trans*-spliced genes (Supplemental Table S1). The 10 % rate is likely to be too low because the quantification pipeline required the full 22 bp SL to be present at read ends [32]. In contrast, SLIDR takes a much more flexible approach that allows for shorter read tails and thus detects substantially more SL-containing reads and SL *trans*-spliced genes.

#### Trichinella spiralis

The parasite *T. spiralis* possesses a diverse and unusual set of 15 SLs that are encoded by up to 48 genes [31] and are spliced to about 30 % of all 16,380 genes [20]. Three out of these 15 SLs (*Tsp*-SL2, *Tsp*-SL10 and *Tsp*-SL12) are SL2-type SLs specialised for resolving downstream genes in operons [20]. We downloaded genome PRJNA12603.WBPS10 and three RNA-Seq libraries from bioproject PRJNA510020 [20]. SLIDR was run with parameter `-S ’.{20,50}AT{ 4,6}G`’ to accommodate for the smaller distance of the *Sm* binding motif to the splice donor site [31].

SLIDR detected all 15 known SLs and a total of 22 SL RNA genes (Supplemental Table S1), which is an increase over the original 19 SL RNA genes identified from cDNA evidence [31] and suggests that many of the 29 additional copies in the genome may not be functional [31]. The SLs were assembled from up to 46,266 reads and were spliced to up to 6,200 genes, which yields an SL *trans*-splicing rate of at least 38 % (Supplemental Table S1).

#### Trichuris muris

*T. muris* is a gastrointestinal parasite closely related to *Trichinella spiralis* and possesses 13 SLs that, unlike those of *T. spiralis*, resemble *C. elegans* SLs and are encoded by 13 genes [52, 67]. Three of these SLs (*Tmu*-SL1, *Tmu*-SL4 and *Tmu*-SL12) are SL2-type SLs [52]. The genome-wide extent of SL *trans*splicing in this organism is unknown [52]. We downloaded genome assembly PRJEB126.WBPS15 from WormBase and five unstranded 2×100 bp libraries from bioproject PRJEB1054. SLIDR was run with `-S ’.{25,50}AT{4,6}G`’ to account for a shorter distance of the *Sm* binding motif to the splice donor site [52].

SLIDR detected all 13 known SLs from 15–15,046 reads spliced to 17–3,122 genes (Supplemental Table S1). SLIDR identified 20 SL RNA genes (1-3 per SL), suggesting that some of the SLs are encoded by previously unidentified multiple copies [52]. Additionally, at least three novel SLs were identified from 159–5,958 reads spliced to 168–2,559 genes. Overall, more than 50 % of all 14,995 genes received SLs (Supplemental Table S1).

#### Prionchulus punctatus

A limited SL repertoire of *P. punctatus* has been determined using 5-RACE of cDNA and comprises six SLs that show structural similarity with *C. elegans* SL2 [17]. However, since no genome assembly exists, the genomic organisation of SL genes and the extent of SL *trans*-splicing are unknown [17]. Only two RNA-Seq libraries are available and no reference transcriptome assembly exists. We tested the performance of SLIDR using a *de novo* transcriptome assembly obtained from the same libraries. Illumina adapters and poor-quality bases (phred 30) were trimmed using TRIM_GALORE 0.6.4 [58], transcripts were assembled using TRINITY 2.8.5 [68] and clustered at 100 % sequence similarity using CD-HIT 4.8.1 [69]. The final assembly comprised 141,825 transcripts with an N50 of 786 bp (184-16,745 bp) and total transcriptome size of 74.31 Mbp.

We ran SLIDR with a relaxed *Sm* location range (`-S ’.{25,60}AC?T{4,6}G`’) but only discovered two (*Ppu*-SL1 and *Ppu*-SL3) out of six known SLs, supported by only 327/353 reads and spliced to 122/63 genes respectively (Supplemental Table S1). While these results are little more than initial proof-of-concept, it must be noted that the success of this *de novo* strategy depends critically on the presence of SL RNA sequences in the transcriptome data. Since SL RNAs are not polyadenylated, RNA-Seq library preparation protocols that rely on poly(A) selection will not capture SL RNAs, which limits the use of publically available datasets that were not generated with ribosomal depletion protocols [70, 71] or poly(A)-tailing prior to library preparation [17]. Thus, we expect SLIDR to underperform in transcriptome mode unless a high-quality transcriptome is available.

To illustrate this point, we tested SLIDR on *C. elegans* using the curated transcriptome GCF_000002985.6 _WBcel235 (contains *sls-1* and *sls-2* RNAs) and three stranded 2×150 bp RNA-Seq libraries. SLIDR detected SL1, all eleven SL2 variants, and all 28 functional SL RNA genes, consistent with the results obtained with a genome reference (Table 2). However, the inability to confirm splice acceptor sites means that a large number of false positive candidate SLs were reported (154 candidate SLs in total; Supplemental Table S1). We also note that the AGC clustering mode in SLIDR yielded superior coverage in this dataset due to the large number of long read tails present in the dataset.

### SLOPPR performance in nematodes with SL2-type SL specialisation

SLOPPR is the first tool for predicting SL subfunctionalisation and operons from genome-wide SL *trans*splicing events. We have previously used the same strategy that is now implemented as SLOPPR to comprehensively discover operons in the genome of a nematode, *T. spiralis*, for which there had been only limited evidence for operon organisation [20]. Here we examine SLOPPR’s performance in three other nematodes with well-characterised operon repertoires and SL2-type SLs specialised for resolving mRNAs transcribed from the downstream genes of these operons: *C. elegans, C. briggsae* and *P. pacificus*. We also use SLOPPR to confirm whether the nematode *T. muris* possesses SL2-type SLs and provide first insight into the genome-wide landscape of SL *trans*-splicing and operon organisation of this organism [52].

As above for SLIDR, all RNA-Seq data were quality-filtered with TRIM_GALORE 0.6.4 [58]. We ran SLOPPR with default parameters but also explored how relaxed SL2:SL1 cutoffs affect the quality of the predicted operons. For each organism, we examined the overall SL *trans*-splicing rate (numbers of genes receiving SLs given the RNA-Seq data) and identified how many predicted operons matched reference operons (specificity) and how many reference operons were predicted (sensitivity). SLOPPR correctly identified SL1-and SL2-type SL subfunctionalisation in all species and identified up to 96 % of known operons (Table 3, Supplemental Table S2). Predicted operon for all species are available in GFF3 format in Additional File 3.

**Table 3:**
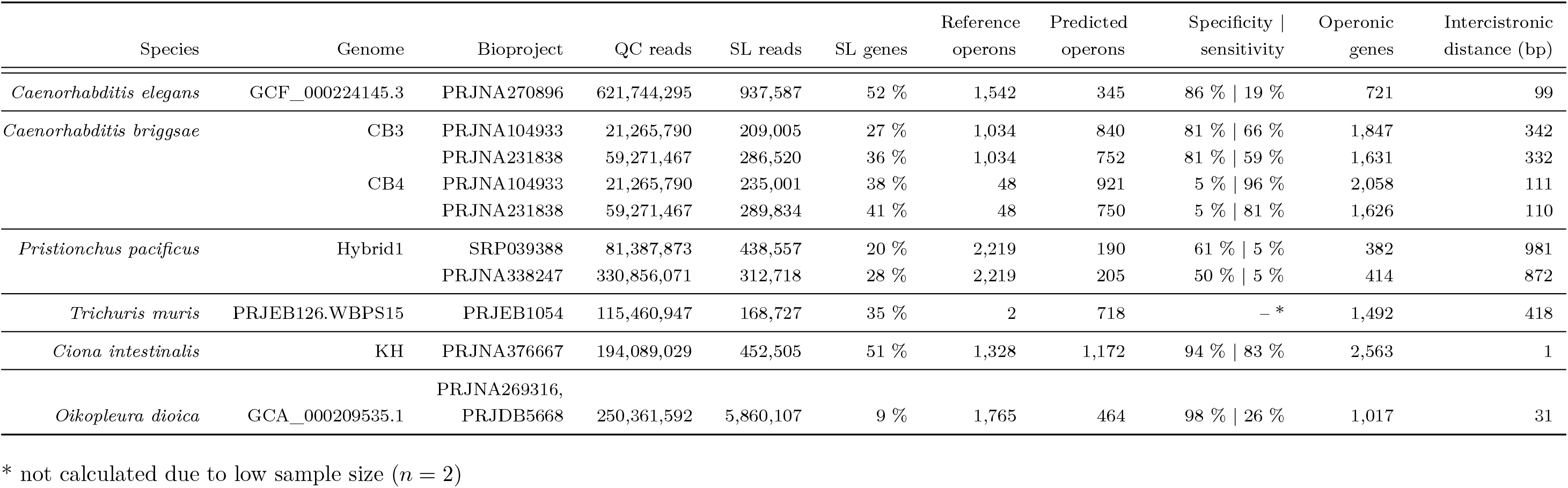
Operon prediction (SLOPPR) results in four nematode and two tunicate species. Identifiers for reference genomes and paired-end RNA-Seq libraries are presented alongside numbers of quality-trimmed reads (QC), numbers of reads with a spliced leader (SL), percentage of genes receiving an SL, numbers of available reference operons, numbers of predicted operons, specificity (predicted operons matching reference operons), sensitivity (fraction of reference operons detected), numbers of operonic genes, and median intercistronic distance among operonic genes. Downstream operonic genes were required to have SL2:SL1 read ratio ≧ 2, and upstream operonic genes were not required to be SL *trans*-spliced. Since the two tunicates *C. intestinalis* and *O. dioica* do not use SL2-type *trans*-splicing, operonic genes were obtained by filtering SL *trans*-spliced genes by intercistronic distance (cutoffs: 84 bp for *C. intestinalis* and 60 bp for *O. dioica*; see main text).

#### Caenorhabditis elegans

Up to 20 % of genes in *C. elegans* are situated in operons, and downstream operonic genes are readily diagnosable by an SL2 bias of 80%–95%, though there are exceptions where downstream genes receive much lower proportions of SL2 [56] or are even SL1-dependent [72]. We designed the operon prediction algorithm in SLOPPR on the basis of SL2:SL1 ratios at genes and benchmarked its performance with a large dataset of 24 unstranded 2×100 bp RNA-Seq runs and 1,542 curated *C. elegans* operons provided with the genome annotations from WormBase (PRJNA13758.WS276). We quantified SL *trans*-splicing events by the canonical SL1 sequence and 11 SL2 variants as supplied by the SL-QUANT pipeline [36].

SLOPPR identified SL *trans*-splicing at 52% of genes, which falls short of the expected SL *trans*-splicing rate of 70-84 % [56]. Of these genes, 36 % were strictly SL1 *trans*-spliced, only 1 % were strictly SL2 *trans*-spliced and 15 % were *trans*-spliced by both SL1 and SL2 (Table 3, Supplemental Table S2). The clustering algorithm correctly identified SL1-and SL2-type subfunctionalisation of SLs (Supplemental Table S2). Using the default SL2:SL1 cutoff of infinity (thus enforcing absence of SL1 at downstream operonic genes), 213 operons were identified, comprising 434 operonic genes with a median intercistronic distance (distance between “gene” GFF annotations) of 105 bp (Supplemental Table S2). Of these operons, 166 (78 %) matched reference operons, but these represented only 11 % of the 1,524 total operons (Supplemental Table S2). We thus relaxed the SL2:SL1 ratio cutoff (`-d 2`), which yielded a final set of 345 operons that comprised 721 operonic genes with 99 bp median intercistronic distance and increased specificity and sensitivity to 86 % and 19 % respectively (Table 3).

While high specificity illustrates that SLOPPR predicts *bona fide* operons and also finds novel candidate operons, the low sensitivity and SL *trans*-splicing rate demonstrates that the 24 RNA-Seq libraries are not nearly large enough to provide exhaustive insight into the SL *trans*-splicing landscape. A meta-analysis of SL *trans*-splicing in *C. elegans* using 1,682 RNA-Seq datasets comprising more than 50 billion reads obtained 287 million reads with evidence of SLs [35]. Even at this large coverage, 97.4% of SL *trans*-splicing events were supported by fewer than 100 reads and a vast number of events with very low read counts could not be distinguished from biological noise in the splicing process [35]. This highlights the inherent limitations of standard RNA-Seq protocols and indicates that, realistically, only a subset of SL *trans*-splicing events and operonic genes can be detected using standard RNA-Seq.

#### Caenorhabditis briggsae

*C. briggsae* is an important comparative model to *C. elegans*, but its gene and operon repertoires are less resolved than those of its relative. The current genome assembly CB4 contains only 48 annotated operons, whereas the older CB3 assembly contains 1,034 operons (of which 51 % were syntenic with *C. elegans*) that were defined based on tight gene clusters that receive SL2 [18]. We decided to examine SLOPPR with both genomes, using the same two unstranded 2×42 bp libraries that were used to define the 1,034 operons in the CB3 assembly [18]. We supplied SLOPPR with the established single SL1 and six SL2 sequences [16].

Using the CB3 assembly, SLOPPR quantified an overall SL *trans*-splicing rate of 27 % and predicted 631 operons comprising 1,346 genes with a median intercistronic distance of 333 bp (Supplemental Table S2). Of these operons, 507 (80 %) matched the 1,034 reference operons (49 % detected). Relaxing the SL2:SL1 ratio cutoff from infinity to two predicted 840 operons, of which 682 (81 %) matched reference operons (66 % detected) (Table 3). Using the CB4 assembly, the SL *trans*-splicing rate was 38 %, and 688 operons were predicted (112 bp median intercistronic distance), of which 37 (5 %) were among the 48 (77 % detected) reference operons (Supplemental Table S2). Relaxing the SL2:SL1 ratio to two (`-d 2`) resulted in 921 operons (111 bp median intercistronic distance) and recovered 46 out of 48 (96 %) reference operons (Table 3).

Although SLOPPR detected the majority of reference operons in both assemblies, two concerns were raised during analysis: First, only 62-69 % of reads aligned to the genome, which may be due to the short read lengths which causes difficulty in aligning these reads across splice sites [18]. Second, the SL *trans*-splicing patterns varied more between the two libraries (L1 vs. mixed life stages) than they did between SL1 and SL2-type SLs, which caused SLOPPR to cluster the SLs by library instead of SL type (Supplemental Table S2). We thus re-ran the analyses with longer reads from five unstranded 2×76 bp libraries, which aligned at much higher rates (83-96 %) and supported higher SL *trans*-splicing rates of 36 % and 41 % for the CB3 and CB4 genome assemblies respectively (Supplemental Table S2). Despite these improvements, SLOPPR predicted fewer operons at equivalent specificity but with somewhat lower sensitivity (at most 81 % of reference operons detected) (Table 3, Supplemental Table S2).

These results suggest that the more recent CB4 assembly has better gene annotations that yield a much lower median intercistronic distance of about 110 bp, which is consistent with *C. elegans* [56]. Resolving the full operon repertoire in this organism will require more effort since the original predictions were based on inferior genome annotations and RNA-Seq data, while the latest better-annotated assembly contains only few curated operons. SLOPPR predicted at least 700 novel operons in the latest assembly, providing a foundation for future curation efforts (Additional File 3).

#### Pristionchus pacificus

*P. pacificus* is another important comparative model to *C. elegans* that resolves operons with SL2-type *trans*-splicing [16]. A comprehensive survey of SL *trans*-splicing events using SL1-and SL2-enriched RNA-Seq data suggested that 90 % of genes are SL *trans*-spliced and that a total of 2,219 operons may exist on the basis of tight gene clusters and SL1/SL2 *trans*-splicing ratios [19]. We used the same SL-enriched unstranded 2×76 bp libraries with the Hybrid1 genome assembly [73] and SNAP genome annotations and operon annotations from http://www.pristionchus.org [19]. We supplied SLOPPR with the two canonical *Ppa*-SL1a and *Ppa*-SL2a sequences that were used for SL enrichment [19].

SLOPPR detected SLs at only 20 % instead of 90 % of genes, even when including the non-enriched library (SRR1182510). The SL1-enriched library (SRR1542610) supported SL1 and SL2 *trans*-splicing at 16.17 % and 0.05 % of genes, consistent with SL1-enrichment. However, the SL2-enriched library (SRR1542630) showed no evidence of SL2-enrichment (0.98 % of genes) but comparable SL1 levels to the non-enriched control library (8.2 % of genes). These results echo the SLIDR results using the same libraries (see above) and would suggest a far lower SL *trans*-splicing rate than 90 % [19]. Due to the low SL *trans*-splicing rate, only 117 operons were predicted, of which 99 (84 %) matched the 6,909 operon-like gene clusters and 67 (57 %) matched the 2,219 plausible reference operons [19] (Supplemental Table S2). Relaxing the SL2:SL1 ratio cutoff (`-d 2`) yielded 190 operons, of which 115 (61 %) were among the 2,219 reference operons (Table 3).

SLOPPR performed similarly with a set of six non-SL-enriched 2×150 bp libraries from bioproject PR-JNA338247, detecting SL *trans*-splicing at 28 % of genes and predicting at most 205 operons, of which 76 (50 %) matched the 2,219 reference operons (Table 3, Supplemental Table S2). Both sets of libraries also yielded similar median intercistronic distances of 785-981 bp (Table 3, Supplemental Table S2). These distances are much larger than the 100 bp expected in *C. elegans* [56] but are consistent with the median distance of 1,149 bp among all 6,909 gene clusters in *P. pacificus* and very poor synteny of these clusters with *C. elegans* (only 37 out of 6,909 clusters are syntenic; [19]). SLOPPR also correctly identified SL1-and SL2-type clusters from genome-wide *trans*-splicing patterns in both library sets, confirming that *Ppa*-SL1a and *Ppa*-SL2a are functionally diverged (Supplemental Table S2).

These observations suggest that curation efforts are required to resolve the operon repertoire of this organism (Additional File 3). SLOPPR produces plausible results given the limitations of relying on RNA-Seq reads covering the 5’ end of transcripts, but overlap with previously predicted operons is relatively poor. These operons were defined by SL1/SL2 *trans*-splicing patterns based on the assumption that all reads from the SL-enriched libraries are from SL *trans*-spliced transcripts [19]. Since only a small fraction of RNA-Seq reads originate from the 5’ end of transcripts, this assumption cannot be confirmed bioinformatically and thus it cannot be ruled out that these libraries contained contaminant non-*trans*-spliced transcripts despite qPCR-based control experiments [19]. Our SLIDR and SLOPPR analyses cast some doubt onto the efficacy of the enrichment and the accuracy of the reference operon annotations.

#### Trichuris muris

*T. muris* is a gastrointestinal parasite of mice and is an important model system for studying mammalian gastrointestinal parasitism. It belongs to the same clade as *T. spiralis* and *P. punctatus* [74]. Comparative work with *T. spiralis* has identified a repertoire of 13 *T. muris* SLs, of which *Tmu*-SL1, *Tmu*-SL4 and *Tmu-* SL12 show structural similarity with *C. elegans* SL2 and are *trans*-spliced to the downstream genes of two operons that are conserved among several nematode species [52]. However, the genome-wide landscape of SL *trans*-splicing and operons in *T. muris* is unresolved [52]. Since only two reference operons have been defined for *T. muris*, we cannot meaningfully validate SLOPPR operon predictions genome-wide, so we focused on testing the hypothesis that the three putative SL2-type SLs show genome-wide SL *trans*-splicing patterns that are distinct from those of the other ten SLs.

We used genome assembly PRJEB126.WBPS15 from WormBase and five unstranded 2×100 bp libraries from bioproject PRJEB1054. We supplied SLOPPR with the 13 known SLs and designated *Tmu*-SL1, *Tmu*-SL4 and *Tmu*-SL12 as SL2-type. SLOPPR detected a relatively high SL *trans*-splicing rate of 35 % and clustered the 13 SLs into the expected groups comprising *Tmu*-SL1, *Tmu*-SL4 and *Tmu*-SL12 versus all other SLs (Fig. 3). SLOPPR predicted 600 operons that comprised 1,229 operonic genes with a median intercistronic distance of 517 bp. This is larger than the c. 100 bp in *C. elegans* but is consistent with the observed elevated intercistronic distance among manually curated *Tmu* benchmark operons [52] and also with the considerably elevated intergenic distance among non-operonic genes (5,445 bp compared with about 3,500 bp observed in all other species in this study; Supplemental Table S2). Relaxing the SL2:SL1 cutoff (`-d 2`) yielded 118 additional operons (Table 3). Operons from both SLOPPR runs contained the two reference operons and confirmed that the third gene of the *zgpa-1* /*dif-1* /*aph-1* operon is not SL2-*trans*-spliced, suggesting the presence of a “hybrid” operon with an internal promoter [52].

**Fig. 3:**
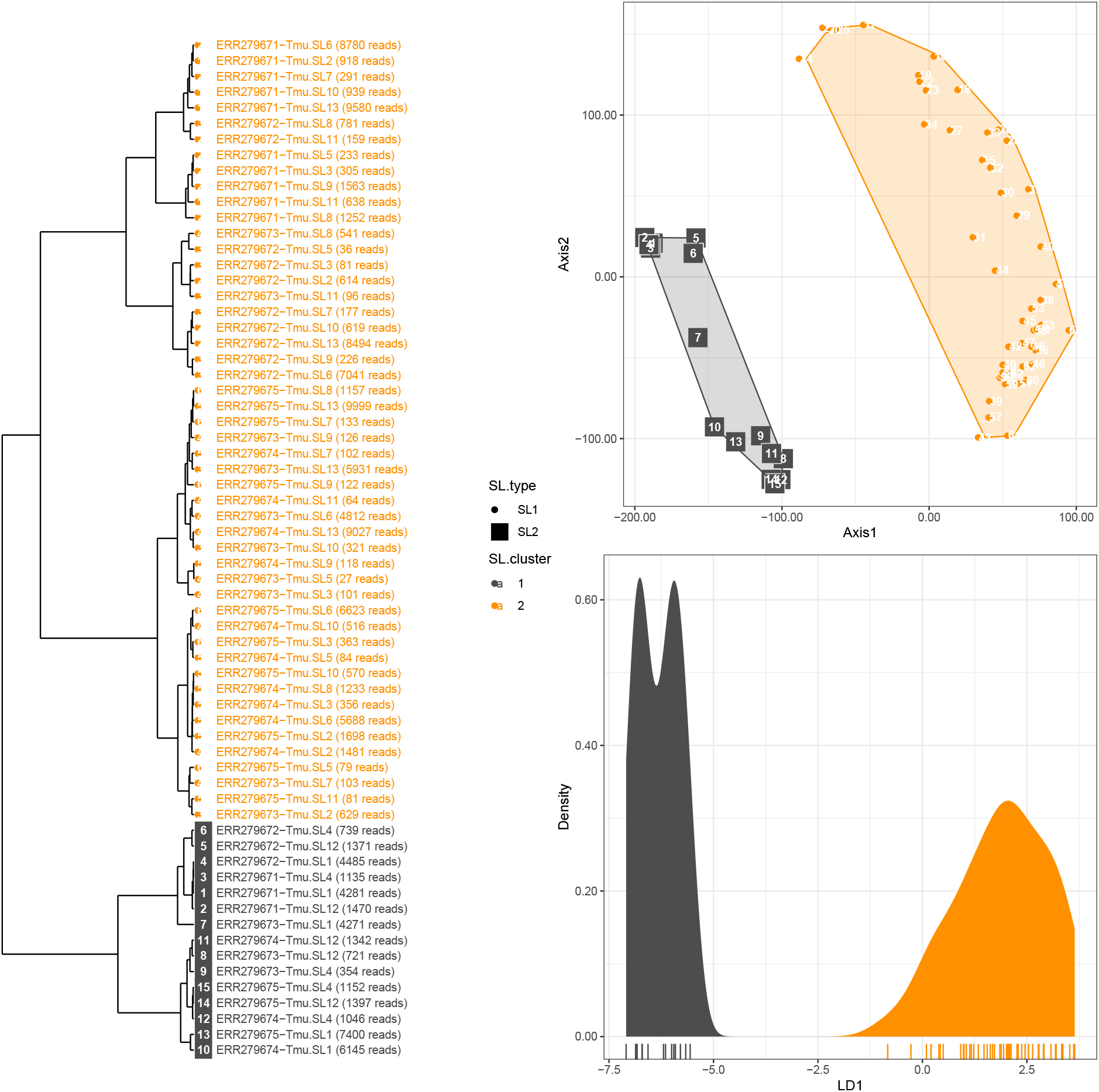
Genome-wide spliced-leader (SL) *trans*-splicing patterns among 13 SLs and five RNA-Seq libraries in *Trichuris muris*. Top right: generalized PCA of normalised genome-wide SL read counts. Symbol shape represents *a priori* SL type (circle: SL1; square: SL2) and colour represents cluster membership inferred via K-means clustering (dark grey: cluster1; orange: cluster2). Numbers inside symbols refer to library identifiers as detailed in the dendrogram on the left (hierarchical Ward’s clustering of PCA eigenvectors). Bottom right: linear discriminant analysis between the two clusters, highlighting complete cluster differentiation by the discriminant function (LD1). *Tmu*-SL1, *Tmu*-SL4 and *Tmu*-SL12 are correctly identified as distinct from all other SLs, confirming their functional specialisation as SL2-type SLs.

These results echo those we obtained in *T. spiralis* [20] and demonstrate that SLOPPR allows for identifying subfunctionalisation among SLs that may correspond to SL1 and SL2-type *trans*-splicing. *Tmu*-SL1, *Tmu*-SL4 and *Tmu*-SL12 are very likely used to resolve polycistronic RNAs in this organism and SLOPPR has predicted plausible candidate operons that warrant curation efforts (Additional File 3).

### SLOPPR performance in the absence of SL specialisation

Having established that SLOPPR accurately predicts operons in organisms that use specialised SLs to resolve downstream operonic genes (SL2-type SLs), we finally aimed to illustrate that SLOPPR is also able to infer operons in organisms that lack such specialisation. Here we demonstrate this ability in two tunicates, *Ciona intestinalis* and *Oikopleura dioica*, both of which possess only a single SL that resolves operons but is also added to monocistronic genes.

In such situations, the SL must be designated as SL2-type such that all genes that receive the SL are classed as operonic; this set of genes will contain *bona fide* operonic genes but will also contain all monocistronic genes that receive the SL. Therefore, these initial candidate operonic genes must be filtered by intercistronic distance to partition out true operonic genes. SLOPPR can be configured to either use a user-supplied cutoff if the expected intercistronic distances are known, or to bisect the distribution of intercistronic distances empirically into two groups using K-means clustering and retaining those genes with short distances. By exploring parameter combinations, specificity and sensitivity in partitioning out operons can be optimised (Fig. 4). Predicted operon for all species are available in GFF3 format in Additional File 3.

**Fig. 4:**
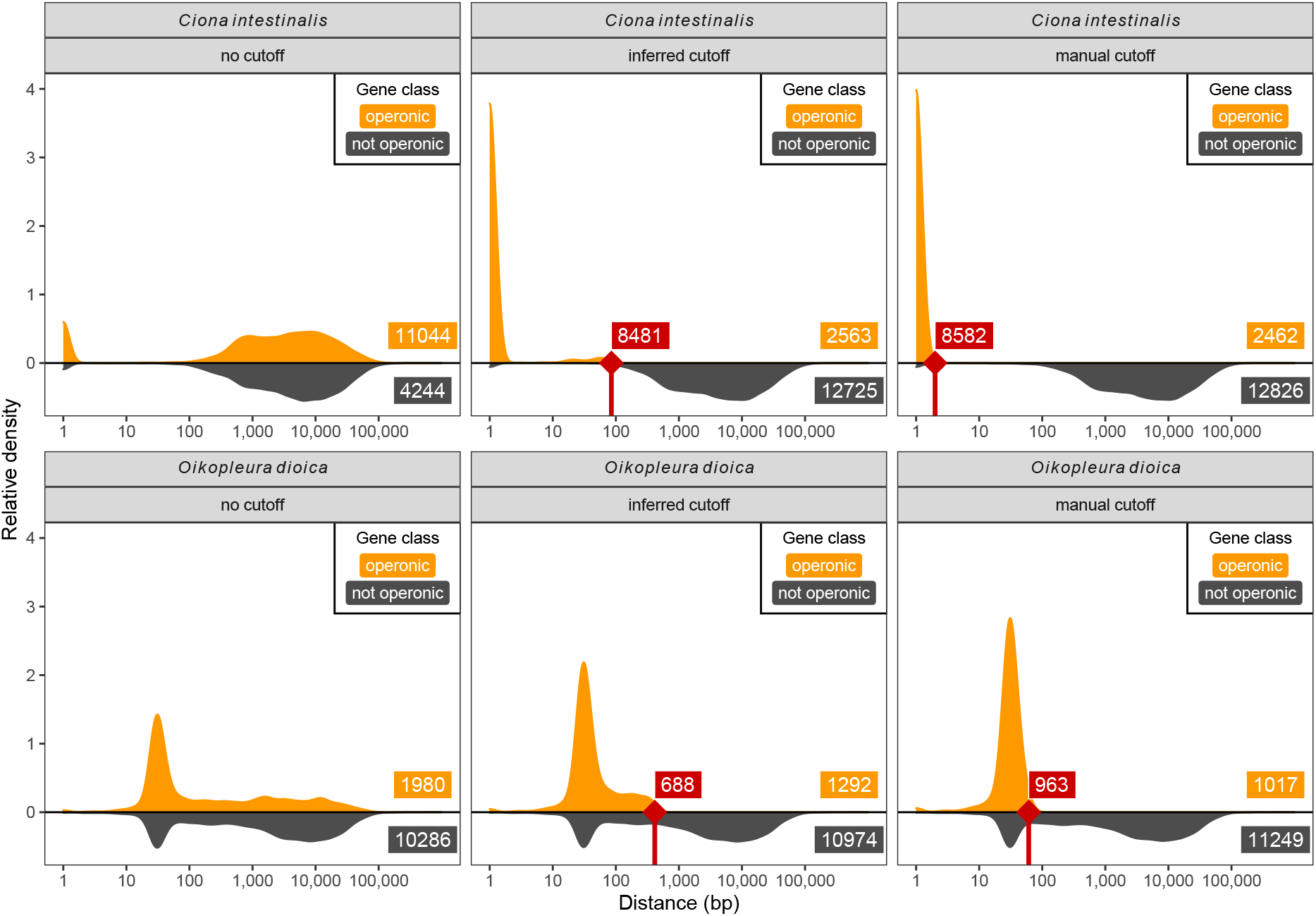
Separation of operonic genes from monocistronic genes in the absence of specialised spliced leaders (SLs), illustrated with SLOPPR data from the tunicates *Ciona intestinalis* (top panels) and *Oikopleura dioica* (bottom panels). All panels display distributions of distances between operonic and non-operonic genes, and labels provide gene numbers. Left panels: in both organisms, a single SL is added to monocistronic and operonic genes, causing SLOPPR to incorrectly designate monocistronic SL-receiving genes with large intergenic distances as operonic. Middle panels: an optimal distance cutoff for operonic genes is inferred via K-means clustering, and genes at or above the cutoff (red notches at 85 and 414 bp respectively) are re-classified as monocistronic non-operonic (red labels). Right panels: a lower manual cutoff (1 bp and 60 bp respectively; red notches at 2 and 61 bp) further reduces the set of genes retained as operonic. Note the peak of tightly-spaced non-operonic genes in the *O. dioica* panels; these genes are likely operonic genes but no SL evidence was obtained from the RNA-Seq data.

#### Ciona intestinalis

The tunicate *C. intestinalis* splices a single SL to downstream operonic genes and infrequently to upstream operonic genes, consistent with operons in nematodes [22, 42]. Using short intergenic distances (<100 bp) as the sole criterion, a total of 1,310 operons comprising 2,909 genes have been predicted [22, 42]. These operons are predominantly dicistronic and have extremely small intercistronic distances, often lacking an intercistronic region altogether [22, 42], similar to the rare SL1-dependent operons observed in *C. elegans* [72]. The genome annotations take SL *trans*-splicing into account and define gene boundaries correctly between poly(A) and *trans*-splicing sites [22].

We obtained the KH genome assembly, the KH gene models (2013) and KH operon annotations (2013; containing 1,328 operons) from the Ghost database (http://ghost.zool.kyoto-u.ac.jp/download_kh.html). We used the same 13 RNA-Seq libraries from bioproject PRJNA376667 that performed best in SLIDR (Supplemental Table S1). SLOPPR detected an overall SL *trans*-splicing rate of 51 %, close to the 58 % expectation [59], despite poor genome alignment rates of 26-64 %. We confirmed with FASTQC 0.11.8 [75] that this was not due to residual adapter contamination or poor sequence quality. Instead, the NCBI SRA Taxonomy Analysis Tool (STAT) indicated that the alignment rates are almost perfectly correlated (Pearson’s *r* = 0.96; *P* = 1*e*^−7^) with the fraction of reads annotated as *C. intestinalis*, suggesting that the libraries are contaminated with other organisms (Supplemental Table S2).

Using the default parameters as for the nematodes, SLOPPR predicted a vastly inflated set of 3,594 operons, of which 1,196 (33 %) matched reference operons and 90 % of the reference operons were detected (Supplemental Table S2). The contamination with SL *trans*-spliced monocistronic genes inflated the intercistronic distances (median 2,287 bp) but a distinct set of genes had very low intercistronic distances, likely representing true operons (Fig. 4). We partitioned out true operonic genes by re-running SLOPPR with automatic inference of the optimal intercistronic distance cutoff (`-i x`) and also requiring SL *trans*-splicing at upstream operonic genes (`-u`). SLOPPR predicted only 856 operons with a median intercistronic distance of 1 bp (inferred cutoff: 68 bp). Of these operons, 823 (96%) matched reference operons, indicating high specificity, but only 62 % of the reference operons were detected (Supplemental Table S2). Re-running the same analysis without requiring SL *trans*-splicing at upstream genes resulted in 1,172 operons, of which 1,100 (94%) matched reference operons and represented 83 % of reference operons (Table 3). The median intercistronic distance was again 1 bp (inferred cutoff: 84 bp), consistent with the notion that many operons in this organism have no intercistronic regions [22, 42].

Finally, to quantify the proportion of operons without intercistronic regions, we re-ran the analysis enforcing a maximum intercistronic distance of 1 bp (`-i 1`). This yielded 1,128 operons, indicating that only 44 operons had intercistronic regions (Supplemental Table S2). Overall, SLOPPR predicted most of the previously proposed operons and also novel operons comprising up to seven genes instead of six as previously reported [22].

#### Oikopleura dioica

Like *C. intestinalis*, the tunicate *O. dioica* possesses only a single SL that is *trans*-spliced to both monocistronic genes and genes in operons, where upstream genes are not required to be SL *trans*-spliced [21, 76]. At least 39 % of genes are SL *trans*-spliced and 58 % of SL *trans*-spliced transcripts originate from operons [76]. A total of 1,765 operons comprising 5,005 genes have been predicted via short intercistronic distances of at most 60 bp [51].

We downloaded genome assembly GCA_000209535.1 (V3) and genome annotations from OikoBase (http://oikoarrays.biology.uiowa.edu/Oiko/), and operon annotations from the Genoscope Oikopleura Genome Browser (https://www.dev.genoscope.cns.fr/oikopleura/). We used four unstranded 2×90 bp libraries from bioproject PRJNA269316 and 16 stranded 2×100 bp libraries from bioproject PRJDB5668, representing various life stages. Similarly to *C. intestinalis*, we observed poor and highly variable background alignment rates (17-69 %) related to poor taxonomic purity (Supplemental Table S2), but recovered large numbers of SL reads (5.8 million in total). However, these reads covered only 9 % of genes, which is much lower than the expected 39 % (Supplemental Table S2).

In default mode, SLOPPR predicted 885 operons with a median intercistronic distance of 57 bp (Supplemental Table S2). Of these, 644 (73 %) matched reference operons. As in *C. intestinalis*, the operons were contaminated with SL *trans*-spliced monocistronic genes having much larger intercistronic distances (median of 2,178 bp) (Fig. 4). Re-running SLOPPR with inference of the optimal intercistronic distance cutoff (`-i x`) yielded 577 operons, of which 521 (90 %) matched reference operons (Supplemental Table S2). The median intercistronic distance was reduced to 33 bp, but the inferred cutoff was still high at 413 bp (Fig. 4). We thus re-ran the analysis with the same hard cutoff of 60 bp (`-i 60`) that was used to predict the 1,765 reference operons [51] and were left with 464 operons (median intercistronic distance of 31 bp), of which 454 (98%) matched reference operons (Table 3).

We also tested the effect of enforcing SL *trans*-splicing at upstream genes (`-u`) across the same three analysis runs and obtained much more stringent sets of 111-165 operons of which 106-143 (87-95 %) matched reference operons (Supplemental Table S2). These results indicate that SLOPPR can discriminate operonic genes from monocistronic genes receiving the same SL, identify the vast majority of previously described operons and also predict a small number of novel operons. One limitation with this dataset was that only few genes received the SL, leading to low sensitivity in detecting known operons (at most 36 %). This was consistent across all libraries tested from several bioprojects and suggests that more RNA-Seq data would need to be generated to fully characterise the SL *trans*-splicing landscape in this organism.

## Conclusions

We have developed two computational pipelines that fill a long-standing gap in our ability to identify and quantify SL *trans*-splicing and eukaryotic operons in any species where RNA-Seq data and reference genomes/transcriptomes are available. SLIDR is a more sensitive, specific and efficient SL discovery pipeline than SLFinder [34], able to uncover a wealth of untapped SL diversity. SLOPPR is the first universal pipeline to predict SL subfunctionalisation and operons from SL *trans*-splicing events, closing this important gap left by existing SL quantification pipelines [35–37]. We have demonstrated here and elsewhere [20] that SLOPPR identifies both *bona fide* and novel operons, blazing the trail for routine operon prediction in any organism with SL *trans*-splicing. Importantly, SLOPPR exploits biological replicates to infer subfunctionalisation among SLs and to moderate noise in SL quantification, which lays a foundational framework for developing a new field of eco-evolutionary “SL-omics”, investigating differential SL usage and *trans*-splicing levels among biological replicates, experimental groups or wild populations.

A fundamental limitation of both SLIDR and SLOPPR is that they were designed for traditional RNA-Seq data where sequencing error is low but only a small fraction of reads originate from the 5’ end of the transcript containing the SL. Most RNA-Seq library preparation methods also show considerable loss of coverage at the 5’ end, which often limits SL detection to a short c. 10 bp portion at typically <1 % of reads [20, 36, 66]. This means that SLOPPR in particular is likely to underestimate the extent of SL *trans*-splicing and operonic gene organisation unless huge amounts of sequencing data are available [35] or specialised SL-enrichment library preparation methods are used [19, 23, 30]. However, our SLIDR analysis on *Hydra vulgaris* vividly demonstrates that SLs at nearly 100 % of all genes can be detected from RNA-Seq data if coverage is sufficient.

We decided to build these pipelines on RNA-Seq data because a wealth of datasets already exists for many species, which continues to grow rapidly. We are thus, for the first time, in the position to investigate SL *trans*-splicing systematically throughout the tree of life without needing to generate novel sequence data. Nevertheless, a powerful future avenue for capturing the full 5’ end of transcripts is direct RNA or cDNA sequencing on the Oxford NanoPore or PacBio long-read platforms [77, 78]. This would require much less sequencing effort because the full molecule is sequenced instead of a short random fraction. SLIDR and SLOPPR could easily be expanded to accept long-read data but would require tailored error-tolerant screening methods to accommodate the higher error rate of NanoPore reads. As these long-read transcriptomics datasets become more commonplace, we expect SL-omics to become a routine molecular tool for uncovering the causes and consequences of this enigmatic source of molecular diversity.

## Supporting information

Operon annotations

## Supplemental material

1. Supplemental Table S1 (.xlsx spreadsheet). Detailed results from all SLIDR runs.
2. Supplemental Table S2 (.xlsx spreadsheet). Detailed results from all SLOPPR runs.
3. Additional File 3 (.zip archive). GFF3 operon annotations generated by SLOPPR.

## Availability and requirements

Project name: SLIDR and SLOPPR

Project home page: https://github.com/wenzelm/slidr-sloppr

Operating system(s): Linux

Programming language: BASH, R

Other requirements: CUTADAPT (tested v2.3), GFFREAD (tested v0.11.4), HISAT2 (tested v2.1.0), BOWTIE2 (tested v2.3.5), SAMTOOLS (tested v1.9), BEDTOOLS (tested v2.28.0), SEQTK (tested v1.3), VSEARCH (tested v2.15.1), BLASTN (tested v2.9.0), SUBREAD (tested v1.6.2), ViennaRNA (tested v2.4.14), R (tested v3.6.0), R packages *data.table, glmpca, ggdendro, MASS, ggplot2, parallel, reshape2*

License: Creative Commons Attribution License (CC BY 4.0)

Any restrictions to use by non-academics: None

## Abbreviations

AGC: abundance-based greedy clustering
CPM: counts per million reads
DGC: distance-based greedy clustering
PCA: principal component analysis
SL: spliced leader
UTR: untranslated region

## Declarations

### Ethics approval and consent to participate

Not applicable

### Consent for publication

Not applicable

### Availability of data and materials

All datasets analysed in this study are publicly available from NCBI (https://www.ncbi.nlm.nih.gov/), ENA (https://www.ebi.ac.uk/ena/browser/home), WormBase (https://wormbase.org/), Ghost database (http://ghost.zool.kyoto-u.ac.jp/download_kh.html), OikoBase (http://oikoarrays.biology.uiowa.edu/Oiko/) and Genoscope (https://www.dev.genoscope.cns.fr/oikopleura/). Accession numbers are detailed in the main text and in supplemental materials. All data generated in this study are included in this published article and its supplemental material.

### Authors’ contributions

JP and BM acquired funding for the project, conceived the research and managed the research activity. MW designed and implemented the computational pipelines with major contributions by JP. MW carried out all data analyses and prepared all display items. MW wrote the manuscript with major contributions by JM and BM. All authors read and approved the final manuscript.

## Acknowledgements

The authors thank Bernadette Connolly for helpful discussions and Andreea Marin, David MacLeod and Lucrezia Piccicacchi for testing the pipelines. The authors acknowledge the support of the Maxwell and MacLeod computer clusters funded by the University of Aberdeen.

